# Acid-sensing ion channel 1a contributes to the prefrontal cortex ischemia-enhanced neuronal activities in the amygdala

**DOI:** 10.1101/2023.08.30.555556

**Authors:** Gyeongah Park, Qian Ge, Zhen Jin, Jianyang Du

**Affiliations:** Department of Anatomy and Neurobiology, University of Tennessee Health Science Center, Memphis, TN 38163, USA; Neuroscience Institute, University of Tennessee Health Science Center, Memphis, TN, United States

**Keywords:** Cerebral ischemia, Stroke, Oxygen-glucose deprivation, Acid-sensing ion channels, ASIC1a, Prefrontal cortex, Amygdala, Endothelin-1, Depression-like behaviors, Synaptic transmission and plasticity, Long-term potentiation

## Abstract

Following a stroke, the emergence of amygdala-related disorders poses a significant challenge, with severe implications for post-stroke mental health, including conditions such as anxiety and depression. These disorders not only hinder post-stroke recovery but also elevate mortality rates. Despite their profound impact, the precise origins of aberrant amygdala function after stroke remain elusive. As a target of reduced brain pH in ischemia, acid-sensing ion channels (ASICs) have been implicated in synaptic transmission after ischemia, hinting at their potential role in reshaping neural circuits following a stroke. This study delves into the intriguing relationship between post-stroke alterations and ASICs, specifically focusing on postsynaptic ASIC1a enhancement in the amygdala following prefrontal cortex (PFC) ischemia induced by endothelin-1 (ET-1) injection. Our findings intriguingly illustrate that mPFC ischemia not only accentuates the PFC to amygdala circuit but also implicates ASIC1a in fostering augmented synaptic plasticity after ischemia. In contrast, the absence of ASIC1a impairs the heightened induction of long-term potentiation (LTP) in the amygdala induced by ischemia. This pivotal research introduces a novel concept with the potential to inaugurate an entirely new avenue of inquiry, thereby significantly enhancing our comprehension of the intricate mechanisms underlying post-stroke neural circuit reconfiguration. Importantly, these revelations hold the promise of paving the way for groundbreaking therapeutic interventions.

## INTRODUCTION

Anxiety disorders and depression occur in 24% and 29-33% of stroke patients, respectively (Hackett et al., 2014), preventing post-stroke rehabilitation and functional outcomes and increasing mortality. Despite the critical consequences, post-stroke anxiety and depression remain poorly understood and treated. Thus, understanding the mechanisms of post-stroke anxiety and depression has important implications for post-stroke outcomes and rehabilitation, and may decrease mortality.

After a stroke, spontaneous recovery occurs through brain remapping (Murphy and Corbett, 2009; Xerri et al., 2014). Newly developed brain circuits lead to different behavioral patterns and new response strategies to recover performance. The neuroplasticity observed in the brain after a stroke might either accelerate recovery or lead to unexpected behavioral changes resulting in emotional symptoms such as anxiety and depression (Naghavi et al., 2019). Although the precise mechanisms by which brain ischemia causes alterations of neuronal activity that result in anxiety- and depression-like behaviors remain poorly understood, we suspect that they most likely involve disrupted neural connections. Neuronal functions in the core and peri-infarct areas as well as the remote brain regions they connect to are disturbed after cerebral ischemia (Murphy and Corbett, 2009). Ischemia-induced infarction in a surface brain region may affect how it projects to the amygdala, which is crucial for the development of anxiety- and depression-like behaviors in mice. Additionally, the increased excitatory neuronal activity in the amygdala is what causes anxiety and depression-like behaviors (Janak and Tye, 2015). The amygdala generally does not experience direct infarction from mild to moderate cerebral ischemia (Cheng et al., 2014), suggesting that the infarction-induced projection to the amygdala may be crucial for the ischemia-induced anxiety- and depression-like behaviors.

During stroke and brain ischemia, reduced blood flow prevents clearance of carbon dioxide, leading to the accumulation of lactic acid in the ischemic brain (Rehncrona, 1985; Siesjo et al., 1996). Brain pH in the infarction sites typically decreases to 6.5-6.0 during ischemia and can be reduced to below 6.0 during severe ischemia (Rehncrona, 1985; Nedergaard et al., 1991; Siesjo et al., 1996). This ischemic acidosis can lead to neurological deficits (Siesjo, 1988; Nedergaard et al., 1991; Choi, 1995; Siesjo et al., 1996). In contrast, protons (H^+^) can also function as neurotransmitters that signal through postsynaptic receptors called ASICs. ASICs play key roles in neurotransmission and synaptic plasticity in the amygdala, a critical site for the formation of emotional behaviors as well as depression and anxiety (Wemmie et al., 2004; Coryell et al., 2007; Coryell et al., 2009; Du et al., 2014). Six ASIC proteins have been identified (ASIC1a, ASIC1b, ASIC2a, ASIC2b, ASIC3, and ASIC4). They are members of the degenerin/ epithelial sodium (Na^+^) channel family. ASICs assemble as homo- or hetero-trimers to form H^+^-gated, voltage-insensitive, Na^+^ and calcium (Ca^2+^) permeable channels that are activated by extracellular H^+^ (Waldmann et al., 1997; Waldmann and Lazdunski, 1998). As a primary target of reduced brain pH in ischemia, ASIC1a enhances neuronal injury by directly or indirectly increasing cellular Ca^2+^ influx (Xiong et al., 2004). ASICs also contribute to amygdala-dependent anxiety- and depression-like behaviors (Coryell et al., 2007; Coryell et al., 2009), which are commonly observed after stroke.

Under physiological conditions, two major counteracting processes control brain pH. Briefly, the utilization of glucose in neuron and glia metabolism generates CO_2_, and/or lactic acid, which results in an acidic pH shift. Counteracting the acidosis, an enhancement of neural activity causes an increase in local blood flow that facilitates the clearance of CO_2_, leading to local alkaline pH shifts (Chesler, 2003). During stroke and brain ischemia, oxygen depletion in the brain forces the utilization of glucose towards anaerobic glycolysis and the reduced blood flow prevents the clearance of CO_2_. Consequently, the accumulation of lactic acid as a byproduct of glycolysis and H^+^ produced by ATP hydrolysis causes pH to decrease in the ischemic brain (Rehncrona, 1985; Siesjo et al., 1996). Brain pH typically falls to 6.5-6.0 during ischemia and can fall below 6.0 during severe ischemia (Rehncrona, 1985; Nedergaard et al., 1991; Siesjo et al., 1996). Ischemic acidosis can then lead to neurological deficits (Siesjo, 1988; Nedergaard et al., 1991; Choi, 1995; Siesjo et al., 1996). In addition, the rapid dynamics of metabolism and CO_2_ clearance suggest that physiological changes in brain pH may have significant consequences for behavior, learning, and memory (Takmakov et al.; Chesler, 2003). An intriguing idea that has emerged from studies of pH-dependent alterations in excitability is that highly localized pH transients might play a signaling role in neuronal communication. Changes in neuronal excitability and synaptic plasticity may include a significant component mediated by pH shift (Kai Kaila, 1998), and H^+^ may function as a ligand to effect these changes. However, how brain ischemia acidosis alters neuronal circuits and influences synaptic transmission and memory is not known.

Testing these novel concepts is expected to open a new area of inquiry and will have a substantial impact on our understanding of the mechanisms that underlie post-stroke neuronal deficits. Finally, this work will lay the foundation for the development of novel therapies for many post-stroke behavioral deficits.

## MATERIALS AND METHODS

### Mice

For our experiment, we used both male and female mice between 10-14 weeks of age. Mice were derived from a congenic C57BL/6 background including wild-type (WT) and ASIC1a knockout (ASIC1a^-/-^). The C57BL6 mice were ordered from the Jackson Laboratory and the ASIC1a^-/-^ mice were gifted from Drs. Michael Welsh’s laboratories at the University of Iowa. All mice were maintained in our animal facilities. Experimental mice were maintained on a standard 12-hour light-dark cycle and received standard chow and water ad libitum. Animal care and procedures met the National Institutes of Health standards. The University of Tennessee Health Science Center Laboratory Animal Care Unit (Protocol #19-0112) and the University of Toledo Institutional Animal Care and Use Committee (Protocol #108791) approved all procedures.

### ET-1 induced brain ischemia model and chemical infusion

For the cannula placement procedure, mice were anesthetized with isoflurane through an anesthetic vaporizer, secured to the stereotaxic instrument, and a cannula made from a 25-gauge needle was inserted bilaterally into the medial prefrontal cortex (mPFC) (relative to bregma: +2.0 mm anterior-posterior; +0.5 mm mediolateral; -2.4 mm dorsoventral). Dental cement secured the cannula and bone anchor screw in place. Mice recovered for 4-5 days before any subsequent testing was carried out. A 10 μL Hamilton syringe connected to a 30-gauge injector was inserted 1 mm past the cannula tip to inject anisomycin (diluted in 1 μl artificial cerebrospinal fluid (ACSF), pH 7.3) over 5 seconds. The injection sites were mapped post-mortem by sectioning the brain (10 μm coronal) and performing cresyl violet staining.

### Brain slice preparation and patch-clamp recording of amygdala pyramidal neurons

Ten minutes after the memory retrieval experiment ended, mice were euthanized with overdosed isoflurane, and whole brains were dissected into pre-oxygenated (5% CO_2_ and 95% O_2_) ice-cold high sucrose dissection solution containing (in mM): 205 sucrose, 5 KCl, 1.25 NaH_2_PO_4_, 5 MgSO_4_, 26 NaHCO_3_, 1 CaCl_2_, and 25 glucose (Du et al., 2017). A vibratome sliced brains coronally into 300 μm sections that were maintained in normal ACSF containing (in mM): 115 NaCl, 2.5 KCl, 2 CaCl_2_, 1 MgCl_2_, 1.25 NaH_2_PO_4_, 11 glucose, 25 NaHCO_3_ bubbled with 95% O_2_ / 5% CO_2_, pH 7.35 at 20°C-22°C. Slices were incubated in the ACSF at least 1 hour before recording. For experiments, individual slices were transferred to a submersion-recording chamber and were continuously perfused with the 5% CO_2_/95% O_2_ solution (∼3.0 ml/min) at room temperature (20°C - 22°C). In some experiments, the regular ACSF was replaced by oxygen-glucose deprivation (OGD, glucose-free) ACSF containing (in mM): 115 NaCl, 2.5 KCl, 2 CaCl_2_, 1 MgCl_2_, 1.25 NaH_2_PO_4_, 25 NaHCO_3_ bubbled with 95% N_2_ / 5% CO_2_, pH 7.35 at 20°C-22°C. The OGD incubation time was indicated in each experiment.

As we described previously (Du et al., 2017), pyramidal neurons in the lateral amygdala were studied using whole-cell patch-clamp recordings. The pipette solution containing (in mM): 135 KSO_3_CH_3_, 5 NaCl, 10 HEPES, 4 MgATP, 0.3 Na_3_GTP, 0.5 K-EGTA (mOsm=290, adjusted to pH 7.25 with KOH). The pipette resistance (measured in the bath solution) was 3-5 MΩ. High-resistance (> 1 GΩ) seals were formed in voltage-clamp mode. Picrotoxin (100 μM) was added to the ACSF throughout the recordings to yield excitatory responses. In AMPAR current rectification experiments, we applied D-APV (100 μM) to block NMDAR-conducted excitatory postsynaptic currents (EPSCs). The peak amplitude of ESPCs was measured to determine current rectification. The amplitude was measured ranging from -80 mV to +60 mV in 20 mV steps. The peak amplitude of EPSCs at -80 mV and +60 mV was measured for the rectification index. In EPSC ratio experiments, neurons were measured at -80 mV to record AMPAR-EPSCs and were measured at +60 mV to record NMDAR-EPSCs. To determine the AMPAR-to-NMDAR ratio, we measured the peak amplitude of ESPCs at -80 mV as AMPAR-currents, and the peak amplitude of EPSCs at +60 mV at 70 ms as NMDAR-currents after onset. For whole-cell LTP recordings, high-frequency stimulation (HFS; 100 Hz, 1 second) was used to induce LTP. For the OGD-induced LTP experiments, the test pulse was delivered regularly during 3 minutes of OGD incubation. The OGD solution will then be rapidly washed out by normal ACSF and continue to record EPSCs for 1-2 hours. The paired acid injection and test pulse were repeated 3 times with a 20-s interval. Data were acquired at 10 kHz using Multiclamp 700B and pClamp 10.1. The mEPSC events (> 5 pA) were analyzed in Clampfit 10. The mEPSC decay time (τ) was fitted to an exponential using Clampfit 10.

### Tail suspension and forced swimming tests

ET-1-induced focal Ischemia and cannula Implantation on Day 0. Prepare an ET-1 solution with a concentration of 2 μg/μl by diluting ET-1 in sterile saline solution (saline solution as the sham control). Ensure proper handling and storage to maintain the solution’s integrity. Anesthetize experimental mice using a VetFlo Isoflurane vaporizer throughout the surgery. Ensure proper depth of anesthesia throughout the procedure. For ET-1 Injection and cannula implantation, use stereotaxic coordinates to guide the injection of ET-1 solution (1 μl each side) into the dual mPFC. Simultaneously, implant a cannula (properly sized for infusion) into the same mPFC region for later drug delivery. For PcTx-1 treatment, behavioral tests on day 14, prepare a PcTx-1 solution with a concentration of 100 nM by diluting PcTx-1 in a sterile saline solution. Prepare a separate saline solution as a control. Administer either PcTx-1 or saline solution into the dual amygdala through the implanted cannula. Allow 90 minutes for the drug to take effect and reach its peak activity. Conduct the tail suspension test to assess depressive-like behavior. Suspend mice by their tails using adhesive tape in a controlled environment. Record and analyze the duration of immobility during a predetermined time period. Conduct the forced swimming test for another group of mice. Place mice individually in a water-filled container and record their swimming and immobility behaviors. Manually analyze the behavioral results by determining the immobility time for each mouse in both the tail suspension and forced swimming tests. Compile the data and calculate the mean immobility time for each experimental group. Perform appropriate statistical analysis (*e*.*g*., ANOVA) to determine significant differences between groups.

### Statistical analysis

One-way ANOVA and Tukey’s post-hoc multiple comparison tests were used for statistical comparison of groups. An unpaired Student’s t-test was used to compare results between the two groups. P<0.05 was considered statistically significant, and we did not exclude potential outliers from our data except the ones that did not receive successful aversive conditioning. The graphing and statistical analysis software GraphPad Prism 8 was used to analyze statistical data, which was presented as means ± SEM. Sample sizes (n) are indicated in the figure legends, and data are reported as biological replicates (data from different mice, and different brain slices). Each group contained tissues pooled from 4-5 mice. Due to variable behavior within groups, we used sample sizes of 10-16 mice per experimental group as we previously described in earlier experiments (Du et al., 2017). In behavioral studies, we typically studied groups with four randomly assigned animals per group, as our recording equipment allowed us to record four separate animal cages simultaneously. The experiments were repeated with another set of four animals until we reached the target number of experimental mice per group. Experimentation groups were repeated in this manner so that each animal had the same controlled environment same time of day and with similar handling, habituation, and processes.

## RESULTS

### OGD (*in vitro* ischemia) enhances ASIC currents in the amygdala

To evaluate whether the expression and function of ASICs are correlated with ischemic conditions in the amygdala, we tested the ASIC currents under an *in vitro* ischemic stroke condition. OGD, a widely used *in vitro* model of ischemia has been shown to increase ASIC currents in cultured cortical neurons (Xiong et al., 2004). Thus, we examined whether OGD affects ASICs in neurons in the amygdala slices. Amygdala slices were perfused with normal ACSF (bubbled with 5% CO_2_ and 95% O_2_) or OGD ACSF (bubbled with 5% CO_2_ and 95% N_2_) for 1 hour and then returned to normal ACSF, and the ASIC-like current was activated by pH 6.0 at -70 mV **(Fig. 1A)**. Activation of ASIC currents with pH 6.0 solution was also performed at indicated time points post-OGD **(Fig. 1B)**. Consistent with a previous study, we found that ASIC currents were potentiated after OGD treatment in the amygdala slices **(Fig. 1B, C)**. These data suggested that ASICs might be the main target of brain acidosis in the amygdala after ischemia (Xiong et al., 2004). We also measured the characteristics of the ASICs after OGD. First, we found the OGD condition leftward drifted the ASIC pH dose-dependent curve to more acidification, and the pH_50_ was changed from pHo 6.4 ± 0.07 to pHo 6.2 ± 0.08, p< 0.001, suggesting that the component of the ASICs was changed after OGD **(Fig. 1D)**; Second, we found the ASIC current desensitization time was prolonged after OGD, supporting the above data that the component of ASICs was changed **(Fig. 1E)**. ASIC1a and ASIC2 are the two major ASIC components in the central nervous system, and they form heterozygotes functional ASICs. Previous studies showed that lack of ASIC2, the current displays a prolonged current desensitization time and more acidic pH_50_ (Askwith et al., 2004; Du et al., 2014). We thus concluded that the OGD condition reduces the ASIC2 proteins in the functional ASICs. To further confirm this conclusion, we tested the ASIC current desensitization time in the ASIC2^-/-^ brain slice before and after OGD. Interestingly, no more change in the desensitization time was found in the ASIC2^-/-^ brain slice, supporting the hypothesis that OGD changes the dynamics of the functional ASICs by reducing the ASIC2 percentage **(Fig. 1F)**. The extended inactivation time suggests that total ASIC-conducted ion influx (Na^+^ and Ca^2+^) per unit time was increased in the neurons from ASIC2^-/-^ mice, resulting in enhanced neuronal activity.

**Figure 1.**
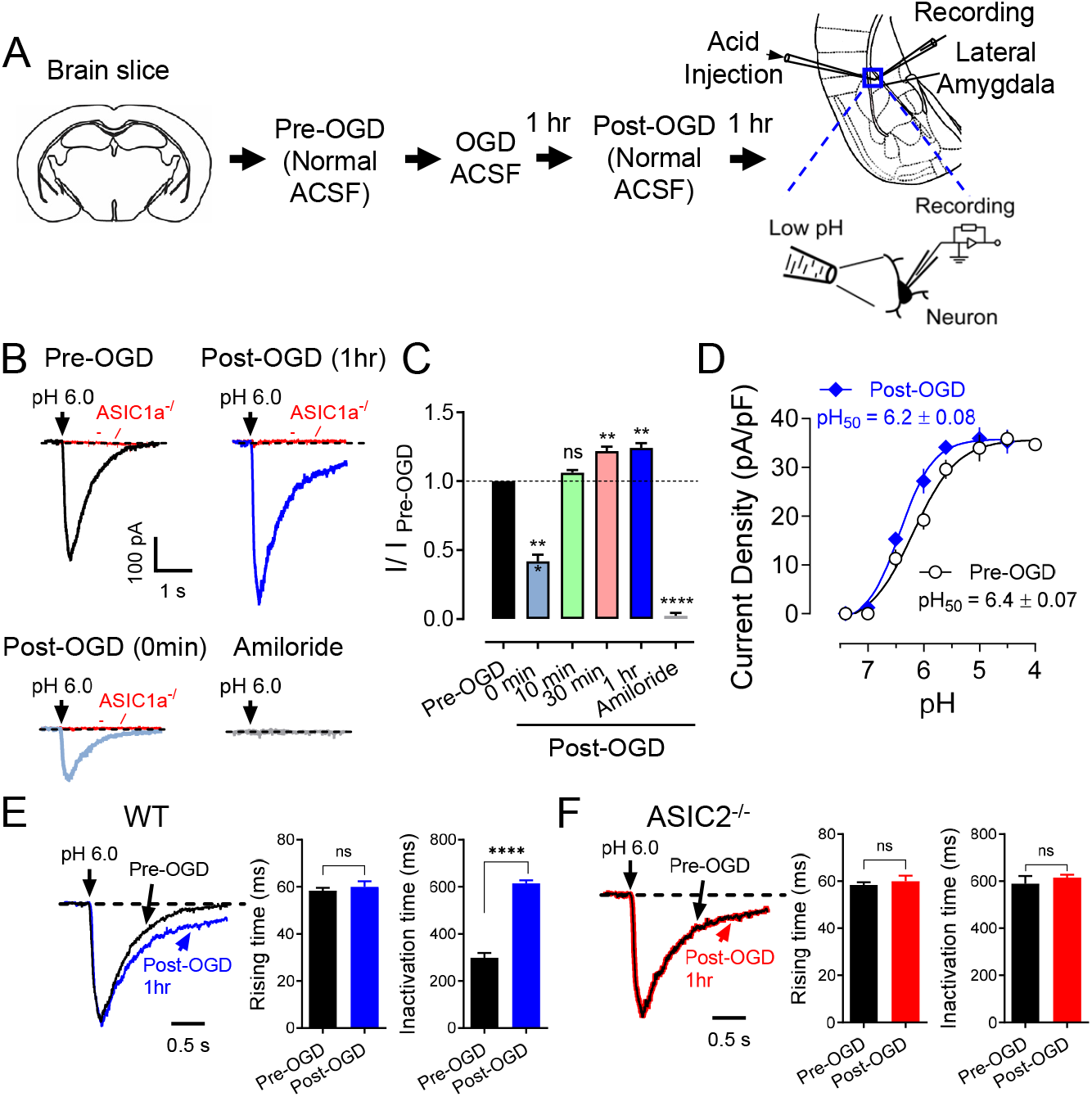
OGD potentiates ASIC1a currents. **(A)** Representative schematic of protocol for the OGD treatment and ASIC current recordings. **(B)** The representative traces of pH 6.0-induced currents before and multiple time points after OGD treatment. **(C)** The summarized data of pH 6.0-induced currents before and after OGD treatment is shown in B. **(D)** pH-dependent activation of ASIC currents before and after OGD. Best-fit yielded pH_50_ of 6.4 ± 0.07 (pre-OGD) and 6.2 ± 0.08 (post-OGD), n = 16-22 cells in 4 mice. **(E)** Normalized acid-induced currents in amygdala slices before and after OGD (black and blue). The decay times in the post-DGD group are longer than in the pre-DGD group, while the rising times remain constant, n = 19 cells in 4 mice. **(F)** Normalized acid-induced currents in ASIC2^-/-^ amygdala slices before and after OGD (black and red), n = 19 cells in 4 mice. ‘n.s.’ indicates not statistically significant. ** p < 0.01, **** p < 0.0001, by unpaired Student’s t-test and one-way ANOVA with Tukey’s post-hoc multiple comparisons. Data are mean ± SEM.

### OGD (*in vitro* ischemia) enhances ASIC1a-dependent synaptic transmission in the amygdala

In addition to our investigation of acid-induced ASIC currents, we embarked on an exploration of ASIC1a-dependent excitatory postsynaptic currents (ASIC1a-EPSCs) in response to ischemic conditions **(Fig. 2A)**. Building upon our previous data that highlighted H^+^ as an activator of ASIC1a-EPSCs in the amygdala, our findings introduced the intriguing possibility that H^+^ and ASIC1a receptors could potentially serve as a neurotransmitter-receptor pair regulating critical neuronal communication. Guided by this notion, we formulated the hypothesis that transient OGD might bolster the strength of ASIC1a-EPSCs. To evaluate this hypothesis, we employed a strategy involving the inhibition of AMPA receptor (AMPAR), NMDA receptor (NMDAR), and GABA receptor (GABAR)-dependent EPSCs using CNQX, D-APV, and PTX cocktails **(Fig. 2B)**. Our results demonstrated a noteworthy enhancement of EPSCs following a mere 10 minutes of OGD, further corroborated by the inhibitory effects of 200 μM amiloride, underscoring the ASIC1a-dependency of these alterations **(Fig. 2B, C)**. Remarkably, we observed that the potentiation of ASIC1a-EPSCs induced by OGD exhibited a time-dependent characteristic, with peak enhancement occurring approximately 30 minutes post-OGD **(Fig. 2B, C)**. This temporal aspect suggests a dynamic process of synaptic modulation in response to ischemic conditions. It also highlights the potential involvement of ASIC1a in mediating the molecular events that lead to this time-dependent strengthening of synaptic transmission.

**Figure 2.**
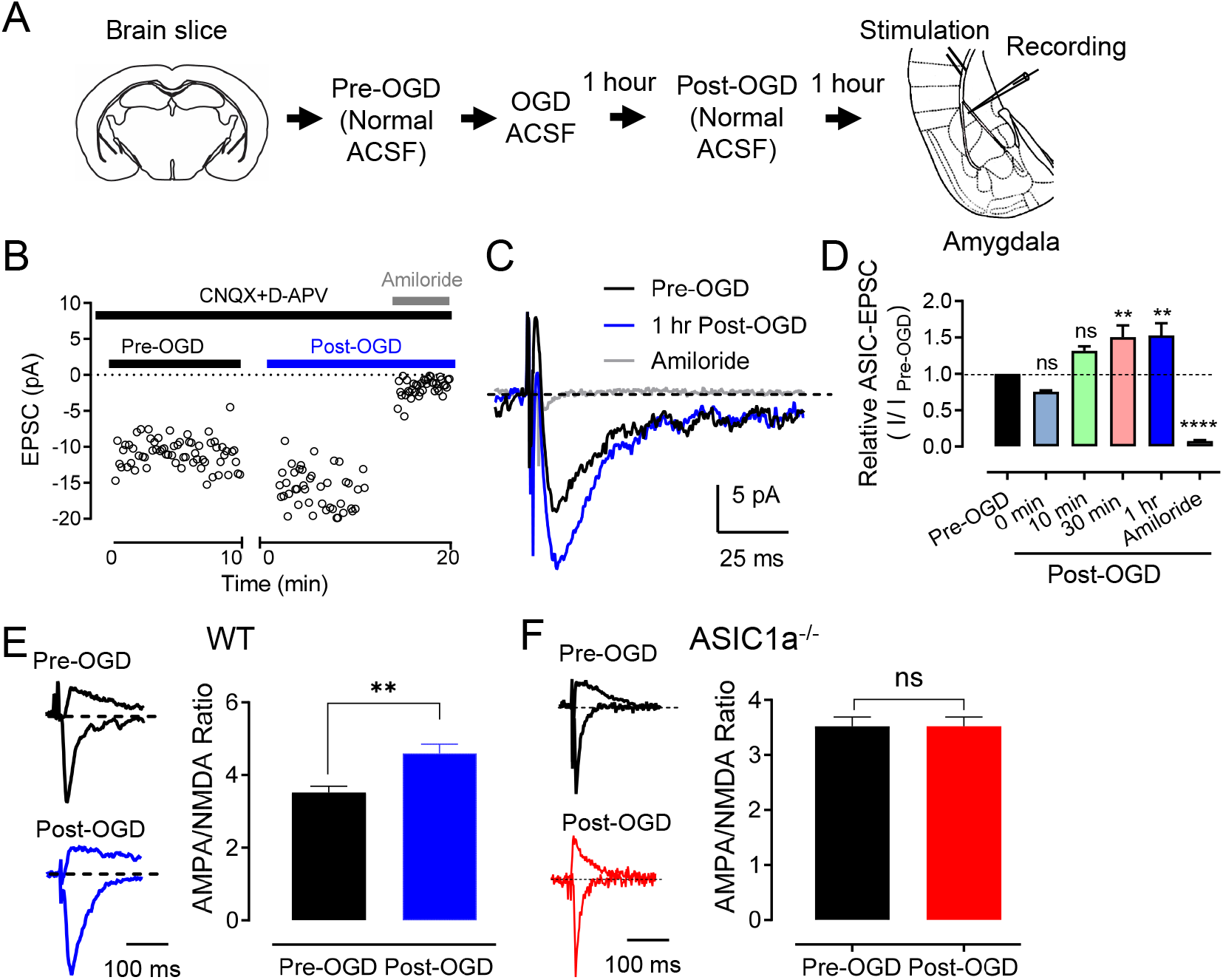
OGD potentiates ASIC1a-dependent EPSCs. **(A)** Representative schematic of protocol for the OGD treatment and EPSC recordings. **(B)** EPSC recordings in amygdala slices. To induce EPSCs, test pulses (100 μs, 0.1 Hz) were delivered through extracellular bipolar electrodes placed on cortical input. Slices were perfused with 25 CNQX, 50 μM D-APV, and 100 μM PTX, with 200 μM amiloride during the times indicated. **(C)** Representative ASIC1a-EPSCs from amygdala slices before and after OGD incubation. **(D)** The summarized data of the time-dependent ASIC1a-EPSCs before and after OGD incubation shown in B. n = 4-8 slices in 4 mice. ‘n.s.’ indicates not statistically significant. ** p < 0.01, **** p < 0.0001, by one-way ANOVA with Tukey’s post-hoc multiple comparisons. **(E)** Left, representative EPSCs were recorded at −80 mV (AMPAR-EPSCs) and +60 mV (NMDAR-EPSCs) in WT amygdala slices. Right, the summarized data of AMPA/NMDA ratios. Current amplitudes were measured 70 ms after onset. n = 15 slices in 4 mice for each group. **(F)** The AMPA/NMDA ratios from ASIC1a^-/-^ amygdala slices. n = 15 slices in 4 mice for each group. ‘n.s.’ indicates not statistically significant. ** p < 0.01, **** p < 0.0001, by unpaired Student’s t-test. Data are mean ± SEM.

Our findings additionally extended into the realm of glutamate receptor (GluR)-dependent EPSCs, which play a central role in synaptic communication. Drawing from previous research indicating that the ratio of AMPA/NMDA EPSCs serves as an indicator of synaptic strength (Rao and Finkbeiner, 2007), we set out to discern how OGD influences this ratio. Encouragingly, our data revealed an augmented AMPA/NMDA ratio in brain slices subjected to OGD incubation, reflecting an overall strengthening of the synapses **(Fig. 2D)**. Importantly, this augmentation was noticeably diminished in ASIC1a^-/-^ slices, providing strong evidence for ASIC1a’s active participation in regulating these OGD-induced changes **(Fig. 2E)**.

The observed enhancement of ASIC1a-EPSCs, the time-dependent potentiation, and the alteration in the AMPA/NMDA ratio collectively imply that OGD-induced changes might recalibrate the efficiency of synaptic communication in the amygdala. Given that ASIC1a is implicated in mediating these synaptic adjustments, the study underscores the pivotal role of this receptor in the context of ischemia-induced alterations in neurotransmission.

### ASIC1a contributes to OGD-induced LTP

Building upon existing scientific literature that has documented the emergence of OGD-induced immediate LTP, indicating the potential contribution of heightened synaptic functionality in the aftermath of pathological conditions such as brain ischemia (Calabresi et al., 2002), we delved deeper into this intriguing phenomenon. Our investigation employed a cohort of WT mice subjected to a concise 3-minute episode of OGD to unravel the dynamics of LTP under these conditions **(Fig. 3A)**. The results yielded a remarkable outcome, revealing the establishment of robust LTP, with the recorded measurements surging to 148 ± 2% of the pre-OGD baseline (n = 10, p < 0.05). This finding underscores the potential for rapid synaptic strengthening in response to OGD, further suggesting a compensatory mechanism at play to counteract the adverse effects of ischemic insult. However, this landscape exhibited significant alterations when examining ASIC1a^-/-^ mice. In this context, the absence of ASIC1a led to the complete annulment of OGD-induced LTP, evident from the recorded values plateauing at 103 ± 5% of the pre-OGD baseline (n = 8, p > 0.05; **Fig. 3B**).

**Figure 3.**
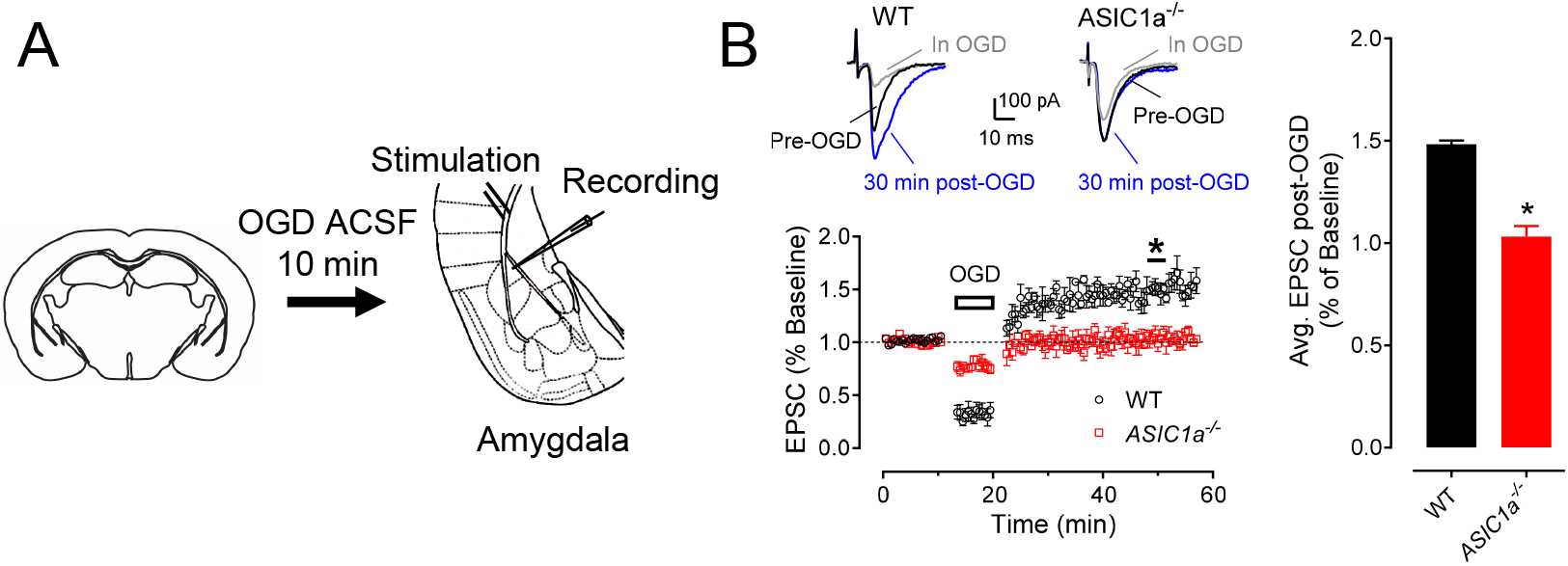
OGD-induced ischemic-LTP in amygdala brain slices. **(A)** LTP after OGD (black trace). Top: representative EPSC traces, pre-OGD, OGD, and 30 minutes after OGD, as indicated. Bottom: EPSCs were recorded in normal ACSF for 10 minutes as a baseline, then the slice was perfused with OGD for 3 minutes and then returned to normal ACSF. Note: EPSCs were not enhanced after OGD in ASIC1a^-/-^ amygdala slices (red trace). **(B)** Summarized data showing relative EPSCs 30 minutes after OGD. n = 4 slices in 4 mice for each group. * p < 0.05, by unpaired Student’s t-test. Data are mean ± SEM.

This striking dichotomy in OGD-induced LTP responses provides a direct glimpse into the pivotal role of ASIC1a in mediating the immediate synaptic potentiation that follows OGD. It highlights ASIC1a as a critical molecular entity orchestrating the rapid adaptive changes in synaptic strength in the wake of ischemic challenges.

### In vivo ischemia enhances ASIC currents and neuronal activities in the amygdala

We employed two distinct in vivo mouse models of ischemia to investigate the effects of focal cerebral ischemia on amygdala neuronal activity and synaptic plasticity. The first model utilized endothelin-1 (ET-1), a vasoconstrictor peptide, to induce focal ischemia within the mPFC (Horie et al., 2008; Chan et al., 2017). A single injection of 1.0 μg of ET-1 or saline (sham group) was administered in the mPFC, with coordinates relative to bregma set at +2.0 mm anterior-posterior, +0.5 mm mediolateral, and -2.4 mm dorsoventral. Subsequently, we explored whether this focal ischemic insult in the mPFC had any impact on Acid-Sensing Ion Channels (ASICs) currents within amygdala neurons. Two days following the ET-1 injection, we sacrificed the animals and prepared amygdala slices for patch-clamp recording **(Fig. 4A)**. The efficacy of the ET-1-induced ischemia in the mPFC was confirmed using crystal violet staining, revealing a distinct ischemic site **(Fig. 4B)**. We then assessed ASIC currents through activation with pH 4.5 solution at specific time points. In alignment with prior research indicating that cerebral ischemia enhances ASIC currents (Xiong et al., 2008), we observed robust potentiation of ASIC-like currents within the amygdala slices subsequent to ET-1-induced ischemia in the PFC **(Fig. 4C)**. These findings suggest the potential involvement of ASIC1a in mediating alterations within the PFC-amygdala circuit under ischemic conditions.

Expanding our investigations, we explored the influence of mPFC ischemia on EPSCs within the amygdala. Following a two-day interval post-ET-1 injection, we sacrificed the animals and conducted evoked EPSC recordings by stimulating cortical inputs and examining miniature EPSCs (mEPSCs) **(Fig. 4D)**. Remarkably, we observed significant potentiation of EPSCs in amygdala slices following ET-1-induced ischemia in the mPFC **(Fig. 4E)**, consistent with previously documented ischemia-induced EPSC enhancements (Quintana et al., 2015). Interestingly, this scenario was notably altered in mice with an ASIC1a deficiency within amygdala slices **(Fig. 4F)**. Similar findings were observed in mEPSC recordings, where we discovered that ET-1-induced ischemia in the mPFC increased mEPSC amplitude in the amygdala **(Fig. 4G)**, however this increase was abrogated in ASIC1a^-/-^ amygdala slices **(Fig. 4H)**.

**Figure 4.**
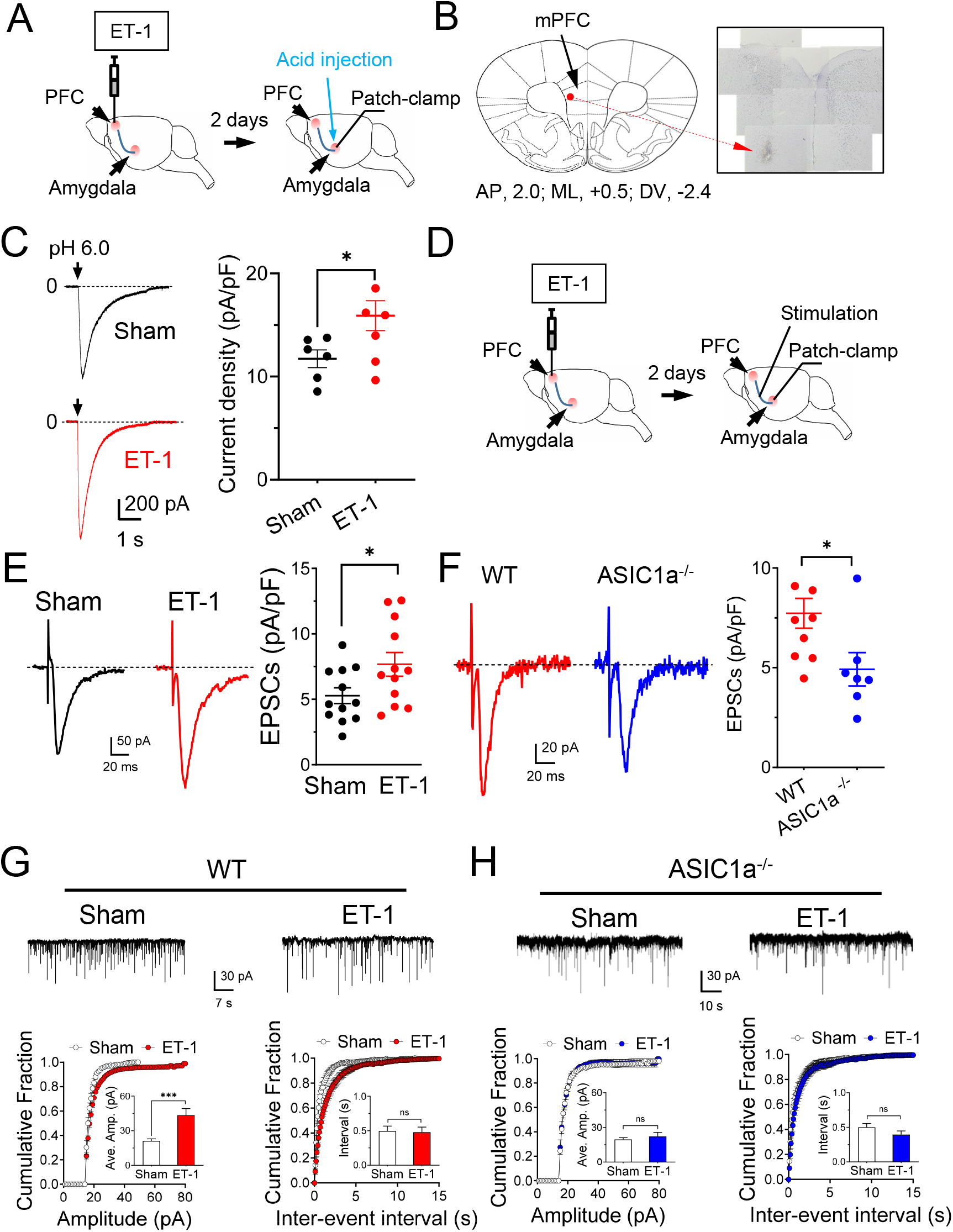
ET-1-induced ischemia in the mPFC potentiates ASIC1a currents and EPSCs in the amygdala. **(A)** Representative schematic of protocol for ET-1 injection and ASIC current recordings. **(B)** Example image of crystal violet staining showing the ET-1-induced ischemia in the mPFC. **(C)** Representative traces (left) and summarized data (right) of pH 6.0-induced currents in the amygdala before and after ET-1 injection in the mPFC. n = 6 cells in 4 mice. **(D)** Representative schematic of protocol for ET-1 injection and EPSC recordings. **(E)** Representative traces (left) and summarized data (right) of EPSCs in the amygdala before and after ET-1 injection in the mPFC. n = 12 cells in 4 mice. **(F)** Representative traces (left) and summarized data (right) of EPSCs in the amygdala after ET-1 injection in the WT and ASIC1a^-/-^ mPFC, respectively. n = 8 cells in 4 mice. **(G)** Upper, representative mEPSC traces in the sham and ET-1 injection groups; lower, cumulative distributions of mEPSC amplitudes and inter-event intervals in the sham (black) and ET-1 injection (red) WT mice. Insets are the summarized data of amplitudes and inter-event intervals. **(H)** The mEPSC results in the sham (black) and ET-1 injection (blue) ASIC1a^-/-^ mice. ‘n.s.’ indicates not statistically significant. ** p < 0.01, **** p < 0.0001, by unpaired Student’s t-test. Data are mean ± SEM.

### The mPFC ischemia-induced depression-like behaviors are ASICs-dependent

Utilizing the ET-1 ischemia mouse model, we established a focused ischemic infarction within the mPFC to investigate emotional behaviors reliant on the amygdala **(Fig. 5A)**. In particular, we explored depression-like behaviors through the implementation of two distinct paradigms **(Fig. 5B)**. The tail suspension test (TST), recognized as an established method for assessing stress and depression in rodents (Castagne et al., 2009), measures immobility as an indicator of helplessness. Notably, our findings revealed an increased duration of immobility in mice injected with ET-1 within the PFC group as compared to the sham group **(Fig. 5C)**. This observation strongly supports the hypothesis that focal ischemia within the PFC exacerbates depressive-like behaviors, with the involvement of the amygdala in this phenomenon. To further scrutinize depressive-like behaviors, we employed the force swimming test (FST), another well-regarded measure of depressive-like tendencies. Injection of the mPFC groups with ET-1 led to a significant elevation in the time of immobility when contrasted with the sham-treated group **(Fig. 5D)**. Collectively, these data indicate that targeted ischemia within the PFC has the capacity to induce a state of depressive-like behavior, offering insight into the complex interactions between brain regions in emotional regulation. To elucidate the role of the ASIC1a subunit in these anxiety and depression-like behaviors, we administered a specific ASIC1a blocker, 100 μM PcTX-1, 90 minutes prior to the commencement of the behavioral tests, encompassing both TST and FST. Intriguingly, our results indicate that the suppression of ASIC1a through PcTX-1 administration yields a substantial reduction in the exacerbated immobility observed in both TST and FST in the context of mPFC ischemia **(Fig. 5C, D)**. This finding lends compelling support to the premise that the manifestation of depression-like behaviors induced by ET-1-driven ischemia within the mPFC is intricately linked to the presence of ASIC1a. By selectively blocking ASIC1a, we observe mitigation of these behaviors, implying a pivotal role for this ion channel subunit in influencing the emotional response to ischemic insults.

**Figure 5.**
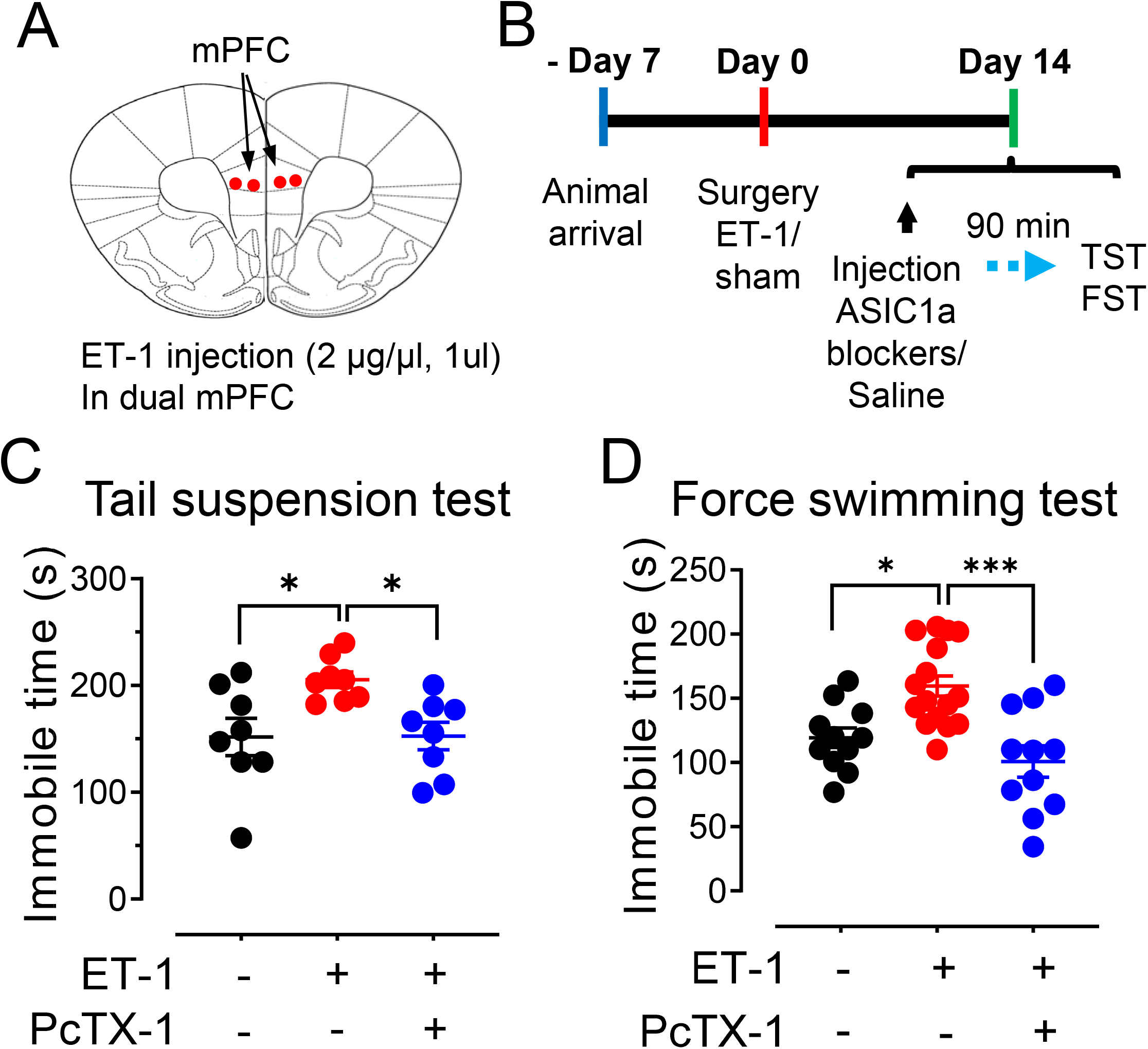
ET-1-induced focal ischemia in the mPFC potentiates depression-like behaviors. **(A)** Representative schematic of protocol for ET-1 injection in dual mPFC. **(B)** Representative schematic of behavioral paradigm. **(C)** The summarized results of the tail suspension test. The injection of 1 μM ET-1 and 100 nM PcTx-1 were indicated in B. n = 8 mice in each group. **(D)** The summarized results of the forced swimming test. n = 11-16 mice in each group. * p < 0.05, *** p < 0.001, by one-way ANOVA with Tukey’s post-hoc multiple comparisons. Data are mean ± SEM.

## DISCUSSION

Understanding the molecular mechanisms that underlie neuron dysfunction after stroke is crucial for developing effective neuroprotective strategies. Cerebral ischemia is a major cause of stroke, which leads to neuronal deficits and brain damage. Dysfunction of synaptic activity is the earliest consequence of cerebral ischemia (Hofmeijer and van Putten, 2012). Elucidating the connections between ischemia and synaptic transmission and plasticity is a key step to understanding and preventing neuron dysfunction after stroke. We hypothesize that ASIC1a is an important therapeutic target that may be modulated to protect against neuronal deficits caused by ischemic stroke and post-stroke mental disorders. The results of our study provide a comprehensive exploration of the impact of in vitro and in vivo ischemic conditions on the function of ASICs and their associated synaptic activity in the amygdala. The role of ASICs in mediating neuronal responses under ischemic conditions has been elucidated, shedding light on their potential contributions to pathological alterations in neural circuits, synaptic plasticity, and behavioral outcomes.

Our investigation into the effect of in vitro ischemia on ASIC currents in the amygdala yielded compelling results. This finding underscores the significance of ASICs in responding to ischemic conditions and suggests that they might be central to the alterations in the PFC-amygdala circuit following ischemic events. Additionally, we observed changes in the characteristics of ASIC currents after OGD, including a shift in the pH dependence and prolonged desensitization time. The results provide strong evidence that ischemic conditions significantly impact ASIC currents in the amygdala. The observed potentiation of ASIC currents following OGD aligns with previous reports in cortical neurons (Xiong et al., 2004; Quintana et al., 2015), indicating a broader role of ASICs in responding to ischemia. The pronounced increase in ASIC currents following OGD suggests that these channels are highly sensitive to ischemic events. This is further supported by the altered pH sensitivity of ASICs after OGD, with the leftward shift of the pH dose-response curve and prolonged desensitization time. These changes point towards a dynamic remodeling of ASIC properties under ischemic conditions, possibly involving alterations in channel subunit composition.

The study delves into the ASIC1a-EPSCs and their modulation by OGD. The enhanced ASIC1a-EPSCs following transient OGD suggest that ASIC-mediated H^+^ signaling plays a role in strengthening synaptic transmission (Quintana et al., 2015). Notably, the time-dependent increase in ASIC1a-EPSCs after OGD highlights a potential mechanism for prolonged synaptic potentiation, which may have implications for information processing and plasticity in the amygdala. The findings related to the AMPAR/NMDAR ratio further underscore the impact of ASICs on glutamate receptor-mediated synaptic transmission. The increase in the AMPAR/NMDAR ratio after OGD suggests enhanced synaptic strength, which aligns with previously established markers of potentiated synapses (Rao and Finkbeiner, 2007). The diminished difference in the AMPAR/NMDAR ratio in ASIC1a^-/-^ slices strengthens the case for ASIC1a’s involvement in these changes. This finding extends our understanding of ASICs beyond their direct H^+^-gated currents to their modulation of glutamate receptor-mediated signaling, indicating a more complex role in synaptic function. The discovery of OGD-induced LTP provides a novel perspective on the interplay between ischemic events and synaptic plasticity. The observed robust LTP following brief OGD exposure suggests that heightened synaptic function under pathological conditions contributes to delayed consequences of brain ischemia, potentially influencing long-term outcomes. Importantly, the significant attenuation of LTP in ASIC1a-deficient mice highlights ASIC1a’s pivotal role in mediating this phenomenon. This finding underscores the significance of ASIC1a in influencing synaptic plasticity dynamics in response to ischemic challenges.

The transition to in vivo ischemia models brings the study closer to the physiological context of brain ischemia. The investigation into ASIC currents and evoked EPSCs following focal ischemia in the mPFC provides evidence for enhanced neural activities in the amygdala. The potentiation of ASIC-like currents following ET-1 injection further emphasizes the role of ASIC1a in mediating the alterations of the PFC-amygdala circuitry in response to ischemic events. This aligns with reports of changes in emotional, memory, and behavioral functions following cerebral ischemia. The implications of the study extend beyond synaptic changes to emotional behaviors. The ET-1-induced focal ischemia in the mPFC results in depression-like behaviors, as evidenced by increased immobility times in the TST and FST tests. The correlation between focal ischemia and depressive behaviors highlights the potential impact of ischemia on emotional regulation. The rescue of these behaviors by the ASIC1a blocker PcTX-1 indicates ASIC1a’s involvement in mediating the effects of ischemia on emotional states.

While this study significantly advances our understanding of ASICs in the context of ischemia, it also raises intriguing questions and potential avenues for future research. Further investigation into the downstream signaling pathways triggered by ASIC activation, the involvement of ASICs in network-wide effects of ischemia, and their role in long-term consequences would provide a more complete picture. Additionally, exploring the potential cross-talk between ASICs and other ion channels and receptors could uncover complex interactions underlying the observed changes.

In conclusion, the study offers a comprehensive exploration of how ischemic conditions influence ASIC1a function, synaptic transmission, and emotional behaviors in the amygdala. The results provide critical insights into the roles of ASICs in mediating synaptic plasticity changes and emotional responses under ischemic challenges. The findings open up new avenues for future research and potential therapeutic interventions targeting ASICs to ameliorate the effects of ischemia on brain function and emotional well-being.

## ACKNOWLEDGMENTS

J.D. is supported by the National Institutes of Mental Health (1R01MH113986) and the University of Tennessee Health Science Center start-up fund.

## AUTHOR CONTRIBUTIONS

J.D. conceived the project. J.D., G.P., Q.G., and Z.J. designed the experiments. J.D. and G.P., performed the patch-clamp experiments and data analysis. Q.G. and Z.J. performed the behavior experiments and data analysis. All authors reviewed and edited the manuscript.

